# The NRF2/ID2 Axis in Vascular Smooth Muscle Cells: Novel Insights into the Interplay between Vascular Calcification and Aging

**DOI:** 10.1101/2024.01.02.573972

**Authors:** Mulin Xu, Xiuxian Wei, Jinli Wang, Yi Li, Yi Huang, Anying Cheng, Fan He, Le Zhang, Cuntai Zhang, Yu Liu

## Abstract

**OBJECTIVE:** Vascular calcification (VC) significantly contributes to cardiovascular morbidity and mortality and escalates with age. However, effective pharmaceutical interventions are lacking, and the molecular mechanisms linking aging to VC remain elusive. This study explores the role of nuclear factor erythroid 2-related factor 2 (NRF2) in VC, specifically focusing on the interplay between vascular senescence and oxidative stress (OS).

**APPROACH AND RESULTS:** Using a chronological aging mouse model, we noted a significant decline in the expression and activity of Nrf2 in the aortas of aged mice, coinciding with increased medial VC. Administering NRF2 activators effectively reduced this calcification. In the vascular smooth muscle cell (VSMC)-specific Nrf2 knockout (*Nrf2^SMCKO^*) models, created using adenine and Vitamin D, there was an increase in calcium deposition within the aortic medial layer, alongside heightened VSMC senescence and OS. Furthermore, NRF2 knockout exacerbated VC in aortic rings and primary VSMCs in high-phosphate conditions, while Nrf2 overexpression in VSMCs inhibited calcium deposition by alleviating cell senescence and OS. RNAseq analysis of the aortas from *Nrf2^SMCKO^* and control mice highlighted a significant downstream regulator of Inhibitor of DNA Binding 2 (ID2). Our results show that reduced NRF2 levels lead to increased VSMC senescence, characterized by heightened p16 expression and diminished ID2 expression. Inhibiting ID2 negated NRF2’s protective effects against VSMC senescence and VC.

**CONCLUSION:** This study emphasizes the critical role of NRF2 dysfunction in the nexus of vascular senescence, OS, and VC. It proposes the NRF2-ID2 axis in VSMCs as a promising therapeutic target for reducing VC and mitigating age-related cardiovascular diseases.

**Highlights:**

- Decline in NRF2 activity is linked to increased vascular calcification (VC) and aging.
- VSMC-specific *Nrf2* knockout causes a remarkable exacerbation of arterial calcification.
- NRF2 alleviates VC by inhibiting oxidative stress and VSMC senescence.
- ID2 contributes to the protective role of NRF2 in VSMC senescence and VC.

## INTRODUCTION

Vascular calcification (VC) in the medial layer of the vessel wall, marked by the ectopic deposition of calcium phosphate crystals, is common in elderly individuals and those with chronic kidney disease or diabetes mellitus^1^. Unlike atherosclerosis, this systemic vascular disorder increases arterial stiffness^2^, leading to diastolic heart failure^3^ and reduced blood flow to critical organs like the brain, kidneys, and liver. Additionally, medial VC can result in chronic limb-threatening ischemia. VC, previously considered a passive degeneration, is now understood as an active process similar to bone formation^4^. The progression of VC hinges on the balance of pro-calcifying and anti-calcifying factors. An imbalance can transform and impair vascular smooth muscle cells (VSMCs), potentially leading to cell senescence, apoptosis, and osteogenic differentiation alongside extracellular matrix (ECM) remodeling^5^. Cellular differentiation is accompanied by cell cycle arrest. A critical view in the late stage of cell senescence is heterotypic differentiation. Oxidative stress (OS) plays a pivotal role in this process, not only by promoting inflammation and DNA damage but also by inducing VSMC senescence^6^. Senescent VSMCs contribute significantly to VC, often displaying an osteoblastic phenotype characterized by elevated levels of BMP-2, RUNX-2, alkaline phosphatase (ALP), and type I collagen^7,8^. This is demonstrated by the elevated p16^Ink4a^ in the calcified arteries of both 5/6-nephrectomized rats and chronic kidney disease (CKD) patients^9^. Therefore, the pro-calcification phenotype of senescent VSMCs is integral to the pathophysiology of VC.

NRF2 (Nuclear factor erythroid 2-related factor 2), a key transcription factor, plays a crucial role in combating oxidative and electrophilic stresses, both exogenous and endogenous. It interacts with ARE/EpRE to regulate the expression of numerous cytoprotective genes, thus preserving the cardiovascular system’s homeostasis^10^. Additionally, NRF2 is involved in bone formation, a process similar to VC, suggesting its potential as a protective agent in this area^11,12^. Various agents, including Empagliflozin, Metformin, Rosmarinic acid, Hydrogen sulfide, and Dimethyl fumarate, have been shown to activate NRF2 effectively, counteracting VC. These agents are noted for mitigating high phosphate-induced calcification in VSMCs and CKD-induced aortic calcium deposition, primarily through antioxidative, anti-inflammatory, anti-apoptotic, and anti-autophagy actions^13–17^. However, it is imperative to acknowledge that NRF2 activators lack specificity and may induce activation of alternative pathways, such as the NF-κB signaling pathway, which has also been implicated in inhibiting VC^18,19^. Interestingly, NRF2 levels are higher in animal models of CKD and in end-stage renal disease (ESRD) patients with VC, even though uremic serum from these patients paradoxically increases VSMC calcification and decreases NRF2^20^. Molecules associated with NRF2, both upstream and downstream, have been identified as inhibitors of VC. For example, glycosylation of KEAP1, which leads to NRF2 degradation, promotes VSMC calcification^21^. Conversely, activating the NRF2/NQO1/HO-1 and NRF2/P62 pathways protects against the osteogenic transition and calcification in VSMCs^22,23^. Nonetheless, NRF2’s effects vary across different tissues and cell types. Its complex and sometimes ambiguous role in whole-body interventions complicates the identification of its primary protective action site. Therefore, creating tissue-specific knockout transgenic models is vital to precisely determine NRF2’s role in vascular smooth muscle tissue, especially in medial VC mechanisms.

Recent research has highlighted the considerable effects of aging on the frequency and intensity of VC, yet the exact mechanisms by which chronological aging influences calcium deposits in blood vessels are still not fully understood^24,25^. An accumulation of senescent VSMCs within aged blood vessels may lead to biological dysfunction and precipitate age-related cardiovascular diseases. A key factor in this process is Prelamin A, predominantly present in VSMCs of older human arteries. It is well-established that Prelamin A interferes with cell division, causes DNA damage, and eventually results in VSMC senescence. Furthermore, it accelerates VSMC calcification by encouraging osteogenic differentiation and the senescence-associated secretory phenotype (SASP)^26,27^. However, a direct link between senescent VSMCs and age-associated VC, as well as a thorough understanding of the regulatory mechanisms involved, is still lacking. Aging is predominantly attributed to an imbalance between oxidant production and antioxidant enzyme activity. Considering the critical role of NRF2 in the antioxidative system, it is hypothesized that NRF2 dysfunction with advancing age could increase reactive oxygen species (ROS) production in the aortas of primates and rodents^28,29^. Significantly, the NRF2 activator, Exendin-4, has shown effectiveness in reducing angiotensin II-induced VSMC senescence^30^. Yet, the connection between VSMC senescence, NRF2 dysfunction, and VC is not fully established. These findings underline the need for more research into the intricate relationship between NRF2 and cell senescence in the context of VC. Such investigations could potentially lead to the development of effective treatments and significant therapeutic breakthroughs.

In our study, we found that aging accelerates VC in a chronological mouse model, characterized by increased OS and reduced apoptosis. Furthermore, we observed a decrease in NRF2 activity in the aortas of aged mice. Interestingly, administering an NRF2 activator was effective in alleviating VC in elderly mice. To further explore this mechanism, we created transgenic *Nrf2* knockout mice specifically in VSMCs (*Nrf2^SMCKO^*) and its control (*Nrf2^WT^*). Results revealed that NRF2 deficiency in VSMCs not only exacerbated VC but also aggravated VSMC senescence and apoptosis. Conversely, overexpressing *Nrf2* in VSMCs mitigated calcification by slowing down cell senescence and apoptosis. This suggests that NRF2’s primary role in counteracting age-associated VC might be to reduce OS and VSMC senescence. Additionally, through RNA-seq analysis comparing *Nrf2^SMCKO^*and control mice, we identified that the protein ID2 (Inhibitor of DNA Binding 2) plays a significant role in NRF2’s protective effects against VSMC senescence and age-related VC. Our findings indicate that the NRF2-ID2 axis in VSMC senescence offers a promising avenue for understanding the connection between aging and VC.

## MATERIALS AND METHODS

### Cell Culture

Primary mouse vascular smooth muscle cells (mVSMCs) were isolated as previously described^31^. Mice were euthanized with intraperitoneal sodium pentobarbital (150 mg/kg), and thoracic aortas from 2-month-old males were harvested. The adventitia and intima were removed, and the aortic media were minced and digested in elastase. Passages 3-8 of primary mVSMCs were utilized. The MOVAS cell line was sourced from ATCC (Rockville, MD, USA). Primary mVSMCs were cultured in Dulbecco’s Modified Eagle Medium/Nutrient Mixture F-12 (DMEM/F-12; Thermo Fisher Scientific) and MOVAS were cultured in Dulbecco’s modified Eagle’s medium (DMEM; Thermo Fisher Scientific), both supplemented with 10% fetal bovine serum (FBS), 100 U/ml penicillin, and 100 mg/ml streptomycin, at 37°C in 5% CO_2_. VSMC calcification was induced in a calcifying medium (DMEM, 10% FBS, 2.6 mM inorganic phosphate) for seven days, with medium changes every two days.

### Animal Studies

Animal protocols were approved by the Institutional Animal Ethics Committee of Huazhong University of Science and Technology (IACUC Number: 3423) and followed NIH guidelines (8^th^ Edition, 2011). We used a Cre/LoxP strategy to generate *Nrf2*-VSMC-specific knockout (*Nrf2*^SMCKO^) mice (**Figure 2A**). *Nrf2^flox/flox^* mice were granted by Pennington Biomedical Research Center/LSU System^32^. Tagln-CreERT2 mice (Cat. NO. NM-KI-225006) were purchased from Shanghai Model Organisms Center, Inc. The F1 generation was backcrossed to obtain *Tagln-Cre^/+^; Nrf2^flox/+^* mice, which were further crossed to generate *Nrf2^SMCKO^* (*Tagln-Cre^/+^; Nrf2^flox/flox^*) and *Nrf2^WT^* (*Tagln-Cre^/-^; Nrf2^flox/flox^*) littermates. In the young group, C57BL/6J male mice aged 2-3 months were included, while in the old group, C57BL/6J male mice aged 18-20 months were included. The animals were maintained on a 12-h light/dark cycle and were given free access to pellets and water.

### Vitamin D-mediated VC mouse model

*Nrf2^SMCKO^* and *Nrf2^WT^* littermates were randomly divided at 8 weeks of age and assigned to either a Vitamin D treatment or its control treatment. In the Vitamin D treatment group, the mice were subcutaneously injected with vitamin D3 (6 or 8 x 10^5^ IU/kg; C1357, Sigma-Aldrich, USA) daily for three days (n=5-8 per group). The vitamin D3 solution (2.64 x 10^6^ IU) was prepared as previously described^33^. Briefly, Vitamin D3 (66 mg) in 200 μl of absolute ethanol was mixed with 1.4 ml of cremophor (S6828, Selleck, USA) and then with 18.4 ml of sterilized water containing 750 mg of dextrose. In the control group, the mice were subcutaneously injected with sterilized water containing dextrose (18.4 ml of sterilized water containing 750 mg of dextrose). After injecting, body weights were monitored daily post-injection until sacrifice.

### Aortic Ring Organ Calcification

Thoracic arteries from *Nrf2^SMCKO^* and *Nrf2^WT^*male mice and young and old C57BL/6J male mice were dissected and cut into 2-3 mm rings after removing the adventitia and intima. These were randomly incubated in either high-phosphate (3.8 mM inorganic phosphate) or regular DMEM at 37°C in 5% CO_2_ for seven days, with medium changes every two days. Calcium deposition was assessed after seven days.

### Chronic Renal Failure Model

The adenine diet-induced Chronic Renal Failure (CRF) mice model was designed following an 8-week program as described previously^34^. Eight-to ten-week-old male mice were used in this model. Adenine was mixed with a casein-based diet to conceal the taste and smell of adenine. Other ingredients of the diet are maize starch (39.3%), casein (20.0%), maltodextrin (14.0%), sucrose (9.2%), maize/corn oil (5%), cellulose (5%), vitamin mix (1.0%), DL-methionine (0.3%) and choline bitartrate (0.2%). Total phosphate content was 0.9% and total calcium content was 0.6%. The control group was fed the same casein diet without the addition of adenine. *Nrf2^SMCKO^* and *Nrf2^WT^* mice were randomly divided into chow and adenine diet groups. All mice were given a 1-week chow diet followed by the 8-week distinct diets (chow vs. adenine), respectively, as shown in **Figure 2C**.

### Quantification of Calcium Content

VSMCs and aortic rings underwent calcification in the calcifying medium. Thoracic aortas from CRF mice were dissected, washed with phosphate-buffered saline (PBS), and incubated with 0.6 N HCl overnight at 4°C. The cell or tissue pellets were dissolved in 0.1 mol/L NaOH and 0.1% SDS for testing the concentration of protein. Calcium content in the supernatant was measured using the QuantiChrom Calcium Assay Kit (C004-2, Nanjing Jiancheng, China) and normalized to overall protein concentration.

### Vonkossa Staining and Alizarin Red S Staining

Artery sections were treated with 5% silver nitrate solution under ultraviolet light for 20-60 minutes; then, the unreacted sliver was removed by incubating 5% sodium thiosulfate for 5 minutes. Nuclei were counterstained with hematoxylin. Calcified nodules appeared brown to black. For Alizarin red S staining, Cells were washed three times with PBS, then fixed with 10% formaldehyde, stained with 2% Alizarin Red S for 30 minutes, and washed with 0.2% acetic acid. Calcification appeared reddish/purple.

### Quantitative Real-Time PCR

Total RNA from aortic tissues and VSMCs was reverse-transcribed to cDNA. The HiScript RT Kit (R222, Vazyme, Nanjing, China) was used according to the instructions as previously described^35^. Primers for mice were listed in the **Supplement Table 1**. Real-time PCR was conducted on a Light Cycler 480 II (Roche, Mannheim, Germany) using ChamQ SYBR qPCR Mix (Q711, Vazyme, Nanjing, China). The delta-delta Ct (2^−ΔΔCt^) method was used for analysis.

### Western Blot Analysis

VSMCs and arterial tissues were homogenized, and protein concentrations were quantified as previously described^35^. Proteins were separated by SDS-PAGE, transferred to PVDF membranes, and immunoblotted with specific primary antibodies: Nrf2 (GTX103322, 1:1000, GeneTex, USA), Nqo1 (11451-1-AP, 1:1000, Proteintech), Ho-1 (ab68477, 1:1000, Abcam, USA), Bmp2 (18933-1-AP, 1:1000, Proteintech, USA), Runx2 (12556, 1:1000, CST, USA), Id2 (3431, 1:400, CST, USA), Caspase-3 (9662, 1:1000, CST, USA), Cleaved Caspase-3 (9664, 1:1000, CST, USA), Tgfb1 (ab92486, 1:1000, Abcam, USA), smad3 (9523T, 1:1000, CST, USA), p-smad3 (AF8315, 1:1000, Affinity, USA), Gapdh (10494-1-AP, 1:1000, Proteintech, USA), β-Tubulin (ABL1030, 1:1000, Abbkine, China). Membranes were incubated with HRP-conjugated secondary antibodies and visualized using an imaging system. Quantification was done using Image J software.

### Histological Analysis

For Immunohistochemistry (IHC) staining, slides were deparaffinized, and endogenous peroxidase activity was quenched with 3% (vol./vol.) hydrogen peroxide for 10 minutes. After blocking nonspecific sites with 10% bovine serum in PBS, slides were incubated with primary and secondary antibodies stained with diaminobenzidine and counterstained with hematoxylin as previously described^35^. The primary antibodies used were listed as follows: Nrf2 (GTX103322, 1:2000, GeneTex, USA), α-smooth muscle actin (α-SMA) (BM0002, 1:400, BOSTER, China), Nqo1 (11451-1-AP, 1:500, Proteintech, USA), Bmp2 (18933-1-AP, 1:100, Proteintech, USA). Positive staining was analyzed using Image-Pro Plus software v 6.0.

Immunofluorescence (IF) staining was performed as described previously^35^. Sections were blocked and incubated with primary and secondary antibodies, followed by DAPI for 5 minutes in the dark. The primary antibodies were α-SMA (BM0002, 1:200, BOSTER, China), Id2 (3431, 1:70, CST, USA), p16 (ab54210, 1:100, Abcam, USA), Nrf2 (GTX103322, 1:500, GeneTex, USA). The fluorescence signal was monitored by confocal laser scanning microscopy (Nikon C2+, Tokyo, Japan).

### siRNA and Lentivirus Transfection

VSMCs were transfected with Small interfering RNA (siRNA) against *Id2* (RiboBio, Guangzhou, China) using Lipofectamine RNAiMAX Reagent (Invitrogen, Carlsbad, CA, USA). The target sequence of mouse siRNA-ID2 was 5’-GACCCAGTATTCGGTTACT -3’. The lentivirus for full-length mouse *Nrf2* (Pubmed No. NM 010902) (LV-NRF2) was purchased from Shanghai Jikai Gene Chemical Technology Co., Ltd and infected MOVAS cells according to the manufacturer’s protocol. A lentivirus carrying green fluorescence protein (LV-GFP) was used as a negative control.

### TUNEL Assay

Aortic tissue slides were processed for *in situ* apoptosis detection using the TUNEL BrightGreen Apoptosis Detection Kit (A112, Vazyme, China). Cells were incubated with permeabilization solution containing Proteinase K (20 μg/ml), and then incubated with a rTdT reaction mixture containing fluorescein-12-dUTP at 3’-OH DNA TdT Enzyme (terminal transferase) following the manufacturer’s instructions. Nuclei were stained with Hoechst. For imaging, a confocal microscopy system was employed. To quantify the extent of apoptosis, six sections with each of ten high-power fields at 400×magnification were averaged. The percentage of TUNEL-positive nuclei was calculated.

### Annexin V-FITC/PI Staining

Cell death was assessed using the Annexin V-FITC/PI Apoptosis Detection Kit (BD Pharmingen, USA). Following the treatments, the cells were collected and washed with cold PBS, and suspended in 195 μL of binding buffer. To stain the cells, 5 μL of Annexin V-FITC and 10 μL of PI (propidium iodide) were added to the suspension, and the mixture was incubated for 20 minutes in the dark at room temperature. After staining, the cells were immediately subjected to flow cytometry analysis to determine the extent of apoptosis.

### γ-H2AX staining assay

Cells were fixed and permeabilized with Triton X-100, then blocked with 1%BSA in PBS for 2 hours. Cells were incubated with primary antibody (γ-H_2_AX, #9718, 1:200, CST, USA) and secondary antibody (AlexaFluor-555, 1:1000, Invitrogen). γ-H_2_AX foci were assessed with red fluorescence. At least three independent experiments were conducted.

### ROS Detection and Superoxide Dismutase (SOD) Activity Measurement

Intracellular ROS levels were determined using dichloro-dihydro-fluorescein diacetate (DCFH-DA) (Beyotime, China). Briefly, after co-culture with t-BHP for 24 hours, the cells were harvested and incubated with DCFH-DA (10 μM) for 20 minutes in the dark at 37 °C. Then, the cells were washed three times with D/F12. The fluorescence signals were recorded using a flow cytometer (ACEA Biosciences, Inc., USA), and representative images were obtained with a fluorescence microscope. Dihydroethidium (DHE, HY-D0079, MedChemExpress, USA) was used to detect ROS production in Optimal Cutting Temperature (OCT)-embedded tissue sections as previously described^35^. SOD activity was measured using a SOD activity assay kit (S0101, Beyotime, China).

### RNA-Seq and Bioinformatics Analyses

RNA-seq libraries from *Nrf2^WT^* and *Nrf2^SMCKO^* mice aortas were prepared using VAHTS mRNA-seq v2 Library Prep Kit for Illumina following the manufacturer’s recommendations, and index codes were added to attribute sequences to each sample. Qubit HS quantification, Agilent 2100 Bioanalyzer/Fragment Analyzer 5300 quality control, the final library size of about 350bp. Analyses RNA-seq libraries were prepared and sequenced on an Illumina NovaSeq platform. Raw data (raw reads) of fastq format were first processed through primary quality control. HTSeq was used to count the read numbers mapped to each gene. Differential expression analysis between two conditions was performed using the DEGSeq R package (version 1.20.0). Differentially expressed genes were defined as those for which the adjusted P-value was below 0.05 and the log^2^ (Fold change) more than 1. GO and KEGG enrichment analysis of differentially expressed gene sets were implemented in the GOseq R and KOBAS 3.0 packages, respectively. GO terms with adjusted P-value below 0.05 were considered as significantly enriched by differentially expressed genes. The Retrieval of Interacting Genes (STRING, http://string.embl.de/) database was used to construct a protein-protein interaction (PPI) network based on the differentially expressed gene sets and the names of all the interacting proteins and the protein-coding genes were extracted from the network. A confidence score of ≥700 was set as the parameter for significance. After obtaining the PPI relationship, a network diagram was constructed with Cytoscape 3.4.0.

### Statistical Analysis

Statistical Analysis Data were analyzed using GraphPad Prism software (version 6.0) and statistical significance was assessed using Student’s t-test and ANOVA. Data are presented as the mean ± SE. All experiments were performed at least 3 independents repeated. Details of the biological replicates are listed in figure legends.

## RESULTS

### Age Exacerbated VC in Mice with Increased VSMC senescence but Decreased Apoptosis

Advanced age is a well-established risk factor for VC. To explore age-related effects on VC, we utilized *ex vivo* aortic ring assays with high phosphate (high-Pi) stimulation, a frequently-used method for studying medial VC. In our study, C57BL/6J male mice aged 19 to 20 months (Aged) and C57BL/6J male mice aged 2 to 3 months (Young) were used to develop an aortic-ring organ culture model. **Figures 1A-B** illustrated that calcification in the aortic ring was significantly induced by 3.8 mmol/L high-Pi stimulation over seven days, as evidenced by von Kossa staining and calcium deposition. Notably, aortic explants from aged mice showed significantly more calcification under high-Pi stimulation compared to those from young mice. Even in the absence of high-Pi stimulation, calcium deposition was marginally higher in the aortic rings of aged mice compared to young ones, although the von Kossa staining showed minimal difference between the two groups without high-Pi intervention. Correspondingly, OS levels were elevated in the aortic rings of aged mice, both with and without high-Pi stimulation, as depicted in **Figure S1A**. IF staining demonstrated increased expression of P16 in aged mice’s aortic rings, particularly under high-Pi conditions (**Figure 1C**). While high-Pi also triggered apoptosis in VSMCs, aged mice’s aortic rings displayed significantly lower VSMC apoptosis under high-Pi conditions compared to younger ones, as shown in **Figures 1D** and **1E**. This suggests that the VSMCs in aged mice’s aortas may be more resistant to apoptosis. Overall, our findings indicate that age exacerbated VC in mice, characterized by increased cell senescence and reduced apoptosis. The evidence points to the possibility that VSMC senescence, rather than apoptosis, contributes to the onset of age-associated vascular calcification.

**Figure 1.**
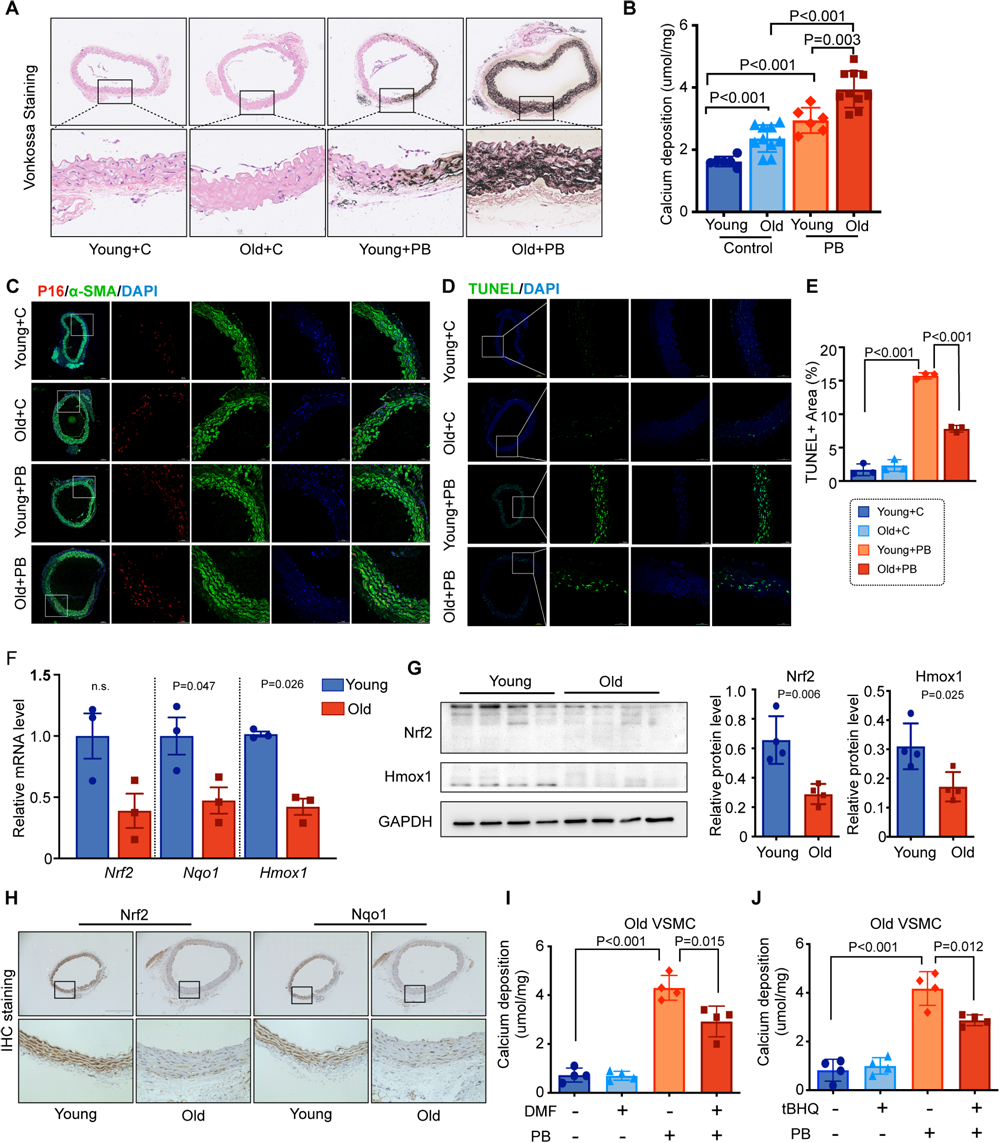
Age Exacerbated VC and Reduced NRF2 Signal in Mice. **A**, Representative micrographs depict von Kossa staining in serial sections of aged and young mice aortas under normal or high-phosphate (High-Pi) conditions. **B,** Quantification of calcium deposition in the aortas of aged and young mice with or without High-Pi stimulation (n=6, 10, 6, and 10, respectively). **C,** Immunofluorescence co-staining illustrates P16 (red) and α-SMA (α-smooth muscle actin; green) in serial sections of aged and young mice aortas under both conditions. **D-E,** Representative micrographs of TUNEL assay and percentage of TUNEL-positive areas in serial sections of aged and young mice aortas, with or without High-Pi stimulation (n=3 per group). **F,** Quantitative real-time PCR results of Nrf2, Nqo1, and Hmox-1 in aged and young mice aortas (n=3 per group). **G,** Western blot analysis of Nrf2 and Hmox-1 in aged and young mice aortas (n=4 per group). **H,** Immunohistochemistry staining displaying Nrf2 and Nqo1 in serial sections of aged and young mice aortas. **I-J,** Calcium deposition quantified in High-Pi induced aged mice aortas with or without NRF2 activator (DMF or tBHQ) intervention (n=4 per group). Scale bars represent 100μm and 50μm. All data are presented as mean ± SE. Statistical significance was determined using a 2-tailed unpaired Student t-test or 1-way ANOVA with Tukey’s multiple comparisons test.

### Reduced NRF2 Signal Observed in the Aged Mouse Aortas

We further explored the impact of aging on NRF2 expression and activity in mouse aortas. Our results showed that while there was no significant difference in Nrf2 mRNA expression between aged and young mouse aortas (**Figure 1F**), NRF2 protein expression was substantially lower in the aortas of aged mice compared to young ones, as demonstrated by western blotting and IHC analysis (**Figures 1G** and **1H**). Additionally, the mRNA levels of NRF2 target genes, including Nqo-1 and Hmox-1, were reduced in aged mice (**Figure 1F**), with a corresponding decrease in protein expression observed by Western blotting and IHC staining (**Figures 1G**s and **1H**). Furthermore, IF assays indicated increased NRF2 nuclear translocation in aged mice aortas compared to young ones under high-Pi conditions (**Figure S1B**), suggesting that aging diminishes NRF2 expression and its functional activity. In addition, the use of NRF2 activators, including DMF and tBHQ, effectively reduced calcification in aged aortic rings under high-Pi stimulation, which was confirmed by observing calcium deposition (**Figures 2I** and **2J**). Therefore, the diminished NRF2 expression and activity in aged mouse aortas could be associated with an increased susceptibility to VC.

### *Nrf2* Knockout in VSMC Aggravated Adenine Diet-induced VC

To explore the relationship between NRF2 and VC, we generated *Nrf2*-VSMC-specific knockout mice (*Nrf2^SMCKO^, Tagln-Cre^/+^; Nrf2^flox/flox^*) by crossbreeding *Nrf2* conditional allele mice (*Nrf2^flox/flox^*) with Cre recombinase mice under the SM22α promoter (*Tagln-Cre^/+^*). Their littermates served as controls (*Nrf2^WT^, Tagln-Cre^/-^; Nrf2^flox/flox^*)(**Figure 2A** and **S2A**). We observed a significant reduction in *Nrf2* expression in the aortas of *Nrf2^SMCKO^*mice compared to *Nrf2^WT^* mice (**Figure 2B**). This reduction in *Nrf2* expression was also noted in the uterus of *Nrf2^SMCKO^* mice, but no significant difference was found in other tissues (heart, liver, and skeletal muscle) when compared with *Nrf2^WT^* mice (**Figure S2B**). Given the high incidence of VC in CRF patients, we utilized an adenine diet-induced CRF mouse model to explore NRF2’s role in VC *in vivo* (**Figure 2C**). The adenine diet induced significant VC in mice, confirmed by von Kossa staining and calcium deposition. Notably, *Nrf2^SMCKO^*mice showed considerably more arterial calcification compared to *Nrf2^WT^* mice in the CRF condition (**Figures 2D** and **2E**). To further substantiate these findings, we conducted additional *ex vivo* aortic ring assays. *Nrf2^WT^* mice aortas exhibited mild VC under high-Pi stimulation. In comparison, aortic rings from *Nrf2^SMCKO^*mice demonstrated a pronounced increase in calcium deposition and von Kossa staining under high-Pi stimulation (**Figures 2F** and **2G**).

**Figure 2.**
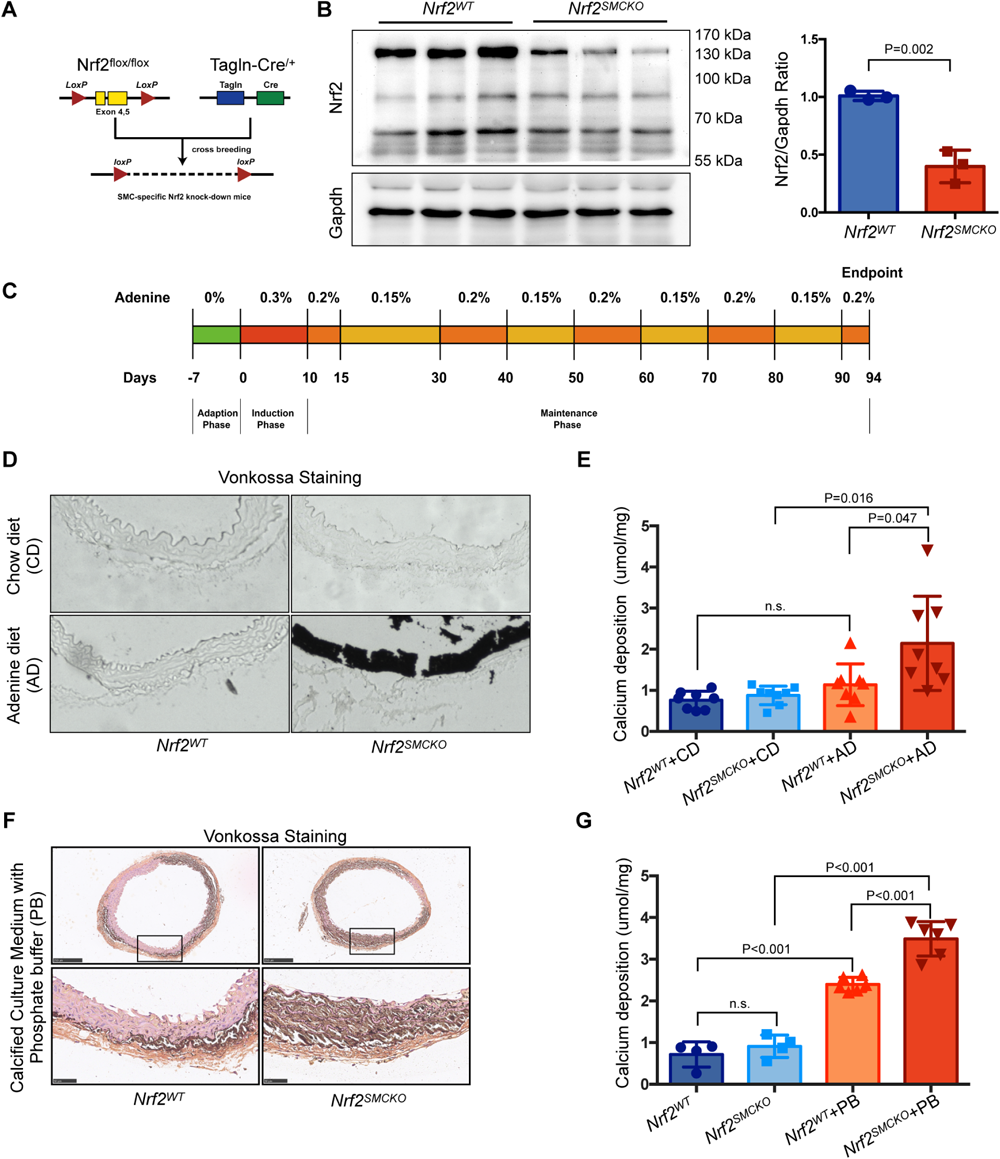
*Nrf2* Knockout in VSMC Aggravated Adenine Diet-induced VC. **A,** Generation strategy for VSMC-specific Nrf2 knockout (*Nrf2^SMCKO^*) mice: LoxP sites flank exon 4 and 5 of the Nrf2 gene. **B,** Western blot analysis of Nrf2 expression in the aortas of *Nrf2^SMCKO^* (*Tagln-Cre^/+^; Nrf2^flox/flox^*) and *Nrf2^WT^* (*Tagln-Cre^/-^; Nrf2^flox/flox^*) littermates (n=3). **C,** Schematic of the adenine diet protocol for inducing chronic renal failure in mice models. **D,** Representative micrographs of von Kossa staining in serial sections of *Nrf2^SMCKO^*and *Nrf2^WT^* mice aortas with chow or adenine diet. **E,** Quantitative analysis of calcium deposition in the aortas of *Nrf2^SMCKO^* and *Nrf2^WT^* mice with chow or adenine diet (n=8 per group). **F,** Representative micrographs of von Kossa staining in serial sections of *Nrf2^SMCKO^*and *Nrf2^WT^* mice aortic rings cultured ex vivo with or without High-Pi treatment. **G,** Quantitative analysis of calcium deposition in *Nrf2^SMCKO^* and *Nrf2^WT^* mice aortic rings cultured ex vivo under High-Pi treatment (n=4, 6, 4, and 6, respectively). Scale bars represent 250μm and 50μm. All data are presented as mean ± SE. Statistical significance was determined using a 2-tailed unpaired Student t-test or 1-way ANOVA with Tukey’s multiple comparisons test.

### NRF2 Deficiency in VSMC Contributes to Vitamin D-mediated VC and Senescence

The Vitamin D-mediated VC mouse model is widely employed to study medial calcification due to its rapid induction and high survival rate ^33^. In our study, both *Nrf2^SMCKO^* and *Nrf2^WT^* mice were subjected to this model to further elucidate the protective role of Nrf2 in medial VC. Nine days post Vitamin D injection, there was a marked increase in Alizarin red S and von Kossa staining in the aortas of both *Nrf2^WT^* and *Nrf2^SMCKO^* mice, with *Nrf2^SMCKO^* mice showing notably higher levels of staining compared to their littermates (**Figures 3A** and **3B**). This was paralleled by a similar trend in calcium deposition (**Figure 3C**). Furthermore, vitamin D treatment significantly induced Bmp2 expression by IHC staining; this effect was intensified in the absence of NRF2, leading to increased BMP2 protein levels (**Figure 3D**). qPCR analysis revealed that Nrf2 knockout increased Bmp2 and Runx2 mRNA levels with vitamin D treatment (**Figure 3E**). Overall, the absence of NRF2 led to a marked increase in VC. Despite vitamin D treatment slightly increasing the mRNA levels of Nrf2 in *Nrf2^SMCKO^*mice, the mRNA levels of Nqo1 had a more obvious increase with vitamin D in both *Nrf2^SMCKO^* and control mice (**Figure 3E**). DHE staining demonstrated that Vitamin D treatment escalated OS in both *Nrf2^SMCKO^* and *Nrf2^WT^* mice, with Nrf2 knockout further amplifying this effect. However, no significant difference in OS levels was observed between the *Nrf2^SMCKO^* and control mice pre-vitamin D treatment (**Figure 3F** and **3H**). This was accompanied by increased VSMC apoptosis in *Nrf2^SMCKO^* mice, as determined by Tunel staining (**Figure 3G** and **3H**). Furthermore, qPCR analysis showed that Vitamin D treatment increased cell senescence markers (p21, p16) and SASP components (Il-6, Il-8, Cxcl1, Csf2, Opg), with these effects being more pronounced in the absence of Nrf2 (**Figure 3I** and **3J**). Even without Vitamin D intervention, Nrf2 deficiency alone elevated cell senescence markers (p21, p16) and certain SASP components (Il-6, Ccl2, Opg) (**Figure 3I** and **3J**). In summary, these results substantiate the hypothesis that NRF2 plays an integral role in mitigating VC by regulating VSMC senescence and apoptosis.

**Figure 3.**
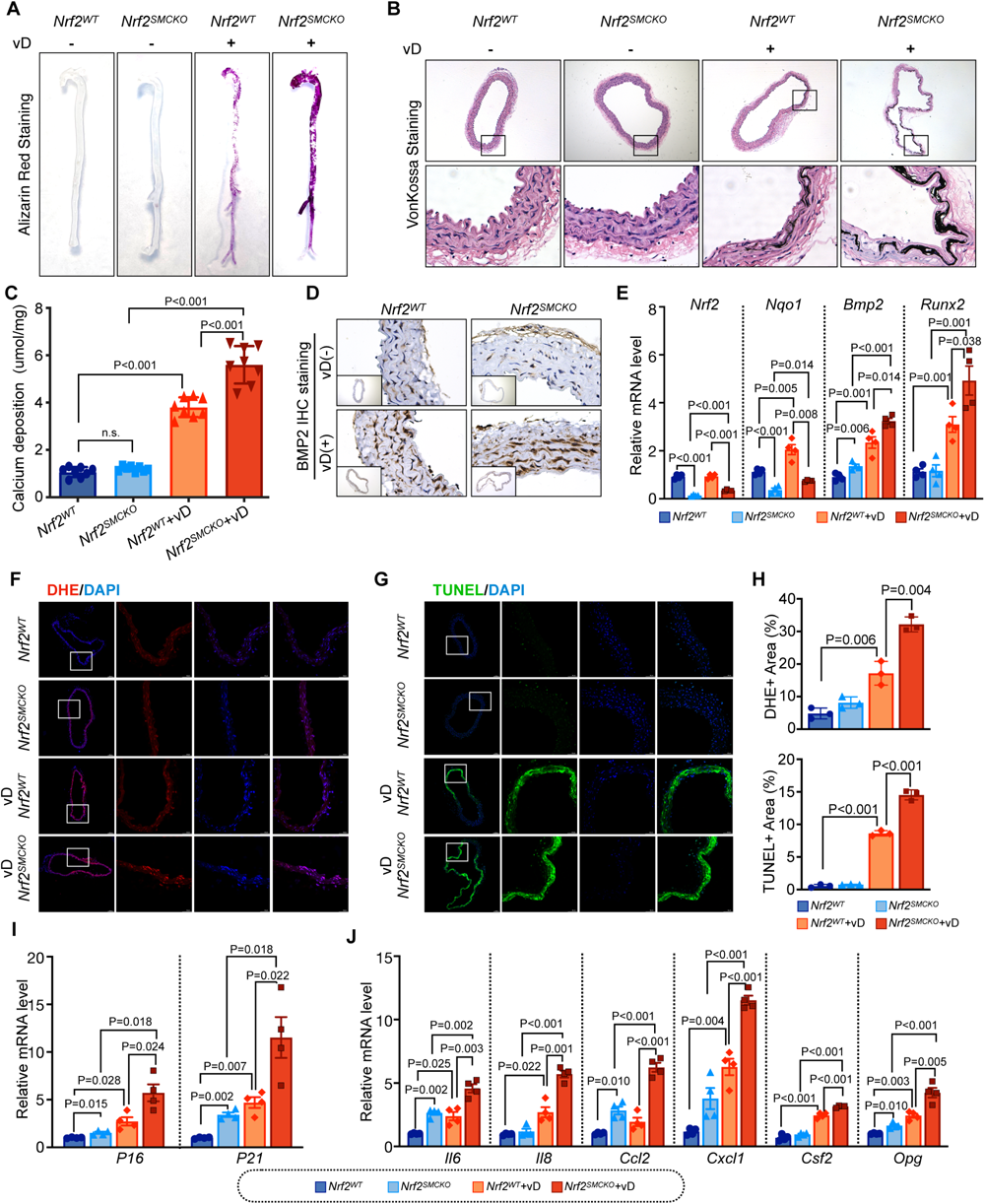
NRF2 Deficiency in VSMC Contributes to Vitamin D-mediated VC and Senescence. **A,** Representative photographs of Alizarin Red S staining in the aortas of *Nrf2^SMCKO^* and *Nrf2^WT^* mice following injection with dextrose water or vitamin D. **B,** Representative photographs of von Kossa staining in serial sections of *Nrf2^SMCKO^*and *Nrf2^WT^* mice aortas post-injection with dextrose water or vitamin D. **C,** Quantitative analysis of calcium deposition in the aortas of *Nrf2^SMCKO^* and *Nrf2^WT^* mice following dextrose water or vitamin D injection (n=8 per group). **D,** Representative photographs of BMP2 immunohistochemistry staining in serial sections of *Nrf2^SMCKO^*and *Nrf2^WT^* mice aortas after dextrose water or vitamin D injection. **E,** Quantitative real-time PCR for Nrf2, Nqo1, Bmp2, and Runx2 in the aortas of *Nrf2^SMCKO^* and *Nrf2^WT^* mice treated with dextrose water or vitamin D (n=4 per group). **F,** Representative micrographs of DHE staining in serial sections of *Nrf2^SMCKO^* and *Nrf2^WT^* mice aortas injected with dextrose water or vitamin D. **G,** Representative micrographs of TUNEL assay in serial sections of *Nrf2^SMCKO^*and *Nrf2^WT^* mice aortas following dextrose water or vitamin D treatment. **H,** Quantitative analysis of the percentage of DHE-positive and TUNEL-positive areas in the aortas of *Nrf2^SMCKO^* and *Nrf2^WT^* mice treated with dextrose water or vitamin D (n=3 per group). **I-J,** Quantitative real-time PCR for p16, p21, and SASP components (Il-6, Il-8, Ccl2, Cxcl1, Csf2, Opg) in the aortas of *Nrf2^SMCKO^*and *Nrf2^WT^* mice following dextrose water or vitamin D injection (n=4 per group). Scale bars represent 100μm and 50μm. All data are presented as mean ± SE. Statistical significance was determined using a 2-tailed unpaired Student t-test or 1-way ANOVA with Tukey’s multiple comparisons test.

### NRF2 Deficiency Irritated VSMC Calcification

To delve deeper into the role of NRF2 in VSMC calcification, we utilized a high-Pi induced VSMC calcification model and evaluated the impact of Nrf2 knockdown on vascular mineralization. We observed a pronounced reduction in Nrf2 and its downstream gene Nqo1 expression in primary VSMCs of *Nrf2^SMCKO^* mice compared to *Nrf2^WT^*mice (**Figure 4A** and **4B**). Despite the NRF2 ablation not exacerbating calcium deposition in VSMCs under normal conditions, primary VSMCs from *Nrf2^SMCKO^* mice exhibited increased calcification susceptibility upon high-Pi exposure (**Figure 4C** and **4D**). To further validate this effect, we employed small interfering RNA (siRNA) to knock down *Nrf2* and examined its direct role in driving VSMC mineralization. While the decrease in Nrf2 expression was substantial at the mRNA level, a concurrent reduction in NQO1 expression was confirmed at both mRNA and protein levels (**Figure S3A** and **S3B**). Aligning with observations in primary VSMCs of *Nrf2^SMCKO^* mice, *Nrf2* silencing alone did not lead to spontaneous calcification without a calcifying medium. However, under high-Pi conditions, *Nrf2* knockdown significantly intensified calcium deposition in VSMCs, contrasting with control siRNA treatments (**Figure S3C** and **S3D**). These *in vitro* findings further substantiate our hypothesis that NRF2 serves as a key inhibitor of VSMC calcification.

**Figure 4.**
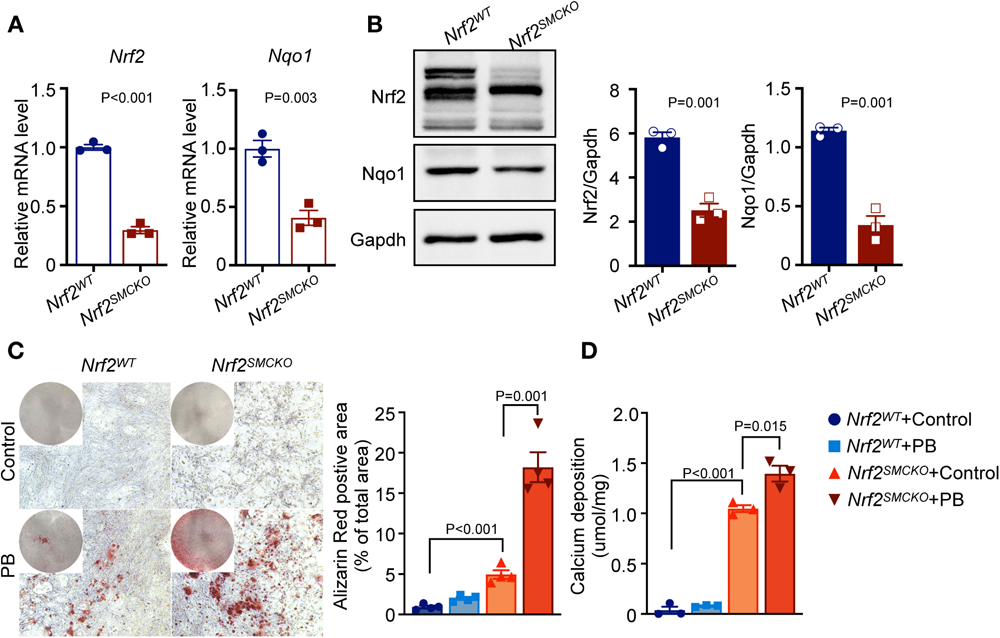
NRF2 Deficiency Irritated VSMC Calcification. **A-B,** Quantitative real-time PCR and Western blot analyses assess Nrf2 and Nqo1 expression in mouse VSMCs derived from the aortas of *Nrf2^SMCKO^*and *Nrf2^WT^* mice (n=3 per group). **C,** Representative graphs of Alizarin Red S staining, both whole well and microscopic, alongside quantification of the percentage of Alizarin Red S-positive area in mouse VSMCs from the aortas of *Nrf2^SMCKO^*and *Nrf2^WT^* mice, with or without High-Pi stimulation (n=4 per group). **D,** Quantitative analysis of calcium deposition in mouse VSMCs derived from the aortas of *Nrf2^SMCKO^* and *Nrf2^WT^* mice under both conditions (n=3 per group). Scale bar represents 20μm. All data are presented as mean ± SE. Statistical significance was determined using a 2-tailed unpaired Student t-test or 1-way ANOVA with Tukey’s multiple comparisons test.

### Nrf2 Overexpression Ameliorated VSMC Calcification

To determine whether Nrf2 overexpression could mitigate VSMC mineralization, we employed lentivirus LV-NRF2 for Nrf2 induction. **Figures 5A** and **5B** illustrated that both mRNA and protein levels of Nrf2 and Nqo1 significantly increased following LV-NRF2 infection compared to LV-Con infection, with negligible variation between conditions with or without high-Pi medium. High-Pi slightly reduced the mRNA levels of Nrf2 following LV-NRF2 infection. However, there is no significant change in NRF2 protein levels. Similarly, Nqo1 expression remained largely unaffected by high-Pi with or without LV-NRF2 infection. Laser confocal microscopy demonstrated a significant enhancement of NRF2 nuclear translocation under high-Pi conditions (**Figure 5C**). Alizarin red staining, indicative of mineralization, was notably intensified in control cells under high-Pi; however, cells infected with LV-NRF2 showed a marked reduction in staining (**Figure 5D**). This reduction was further corroborated by decreased calcium deposition and diminished Bmp2 expression at protein levels, illustrating NRF2’s protective role against VSMC calcification (**Figures 5B**, **5D**, and **5E**). Additionally, γ-H2AX IF staining showed that Nrf2 overexpression mitigated DNA damage in high-Pi conditions (**Figures 5F** and **5G**). Concomitantly, ROS levels significantly decreased, as evidenced by DCFH-DA fluorescent probe (**Figure 5I**) and DHE staining (**Figures 5F** and **5G**), while SOD activity increased in LV-NRF2 infected VSMCs (**Figure 5H**). Flow cytometry using Annexin V-FITC/PI demonstrated that high-Pi markedly escalated total and late apoptosis in VSMCs, both of which were substantially attenuated by Nrf2 overexpression. Contrarily, in the absence of high-Pi, total and early apoptosis rates decreased, and late apoptosis rates were stable in VSMCs with Nrf2 overexpression (**Figure S4A**). Further, both caspase3 and its active form, cleaved caspase3, were reduced with Nrf2 overexpression in high-Pi conditions (**Figure S4B**). Collectively, these results affirm that Nrf2 overexpression effectively counteracts VSMC calcification, diminishing cellular senescence and apoptosis.

**Figure 5.**
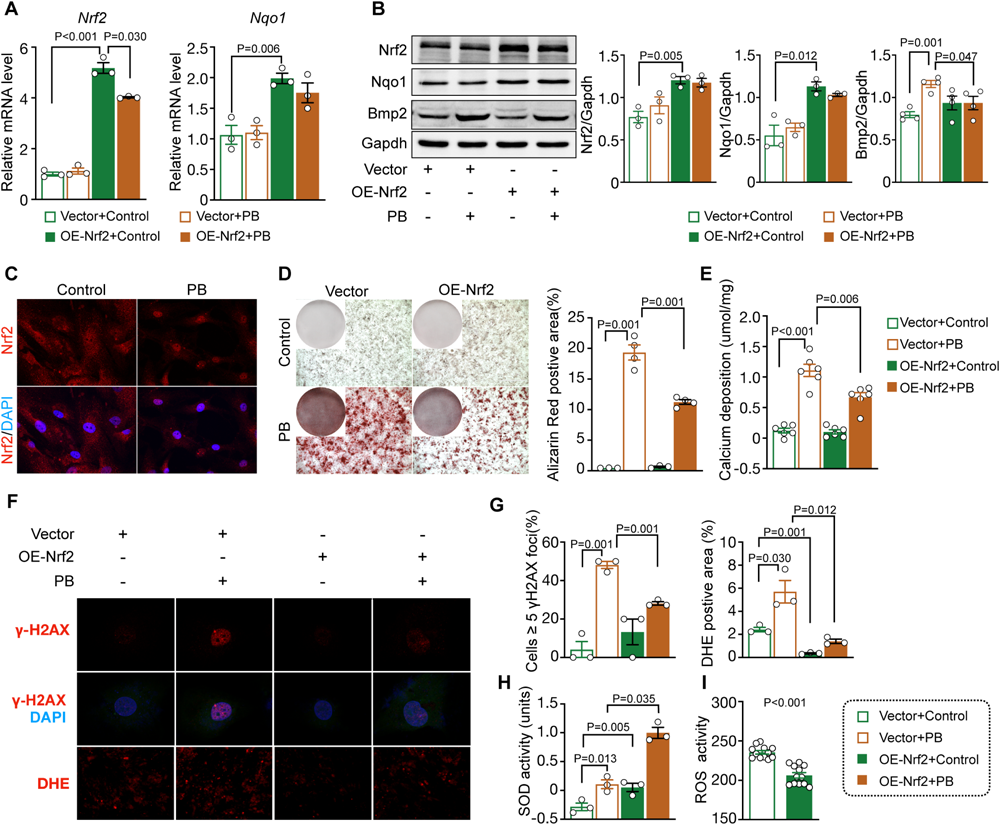
Nrf2 Overexpression Ameliorated VSMC Calcification. **A,** Quantitative real-time PCR analysis of Nrf2 and Nqo1 in MOVAS cells treated with NRF2 lentivirus or control lentivirus, with or without High-Pi stimulation (n=3 per group). **B,** Western blot analysis of NRF2, NQO1, and BMP2 in MOVAS cells treated with NRF2 lentivirus or control lentivirus under both conditions (n=3 per group). **C,** Representative micrographs of NRF2 immunofluorescence staining in MOVAS cells with or without High-Pi stimulation. **D,** Representative graphs of Alizarin Red S staining, both whole well and microscopic, and quantification of the percentage of Alizarin Red S-positive area in MOVAS cells treated with NRF2 lentivirus or control lentivirus under both conditions (n=4 per group). **E,** Quantitative analysis of calcium deposition in MOVAS cells treated with NRF2 lentivirus or control lentivirus, with or without High-Pi stimulation (n=6 per group). **F,** Representative micrographs of γ-H_2_AX and DHE staining in MOVAS cells treated with NRF2 lentivirus or control lentivirus under both conditions. **G,** Quantitative analysis of cells with 5 or more γ-H_2_AX foci and DHE-positive areas in MOVAS cells treated with NRF2 lentivirus or control lentivirus, with or without High-Pi stimulation (n=3 per group). **H,** Measurement of SOD (superoxide dismutase) activity in MOVAS cells treated with NRF2 lentivirus or control lentivirus under both conditions (n=3 per group). **I,** Detection of intracellular ROS (reactive oxygen species) levels using DCFH-DA in MOVAS cells treated with NRF2 lentivirus or control lentivirus, with or without High-Pi stimulation (n=12 per group). Scale bar represents 20μm. All data are presented as mean ± SE. Statistical significance was determined using a 2-tailed unpaired Student t-test or 1-way ANOVA with Tukey’s multiple comparisons test.

### Transcriptional Profiles in *Nrf2*^SMCKO^ Mice Aorta

The dataset comprised 12 samples, including 6 aortas from *Nrf2^WT^*mice and 6 from *Nrf2^SMCKO^* mice. From these, 3477 differentially expressed genes (DEGs) were identified: 2235 were upregulated and 1242 were downregulated (**Supplement Table 2**). Subsequent analysis using DAVID revealed the most significantly enriched Gene Ontology Biological Process (GO-BP) categories and KEGG pathways, as shown in **Figures 6A** and **6B**. Notably, the TGF-β signaling pathway topped the list of enriched KEGG pathways. Further, Gene-set enrichment analysis (GSEA) illustrated that genes in the TGFβ signaling pathway were significantly overrepresented among DEGs in *Nrf2^SMCKO^*mice aortas compared to those in *Nrf2^WT^* mice, suggesting a potential regulatory role of NRF2 in this pathway (**Figures 6C** and **6D**). We then utilized the STRING database to analyze protein-protein interactions (PPIs) among DEGs in the ’TGF-β signaling pathway’, selecting interactions with medium confidence scores (greater than 0.4) for visualization (**Figure 6E**). ID2 emerged as a core gene within the TGFβ signaling pathway and was markedly downregulated among the DEGs. In contrast, P16 was significantly upregulated. The differential expression of both Id2 and p16 was confirmed in the aorta samples of *Nrf2^WT^* and *Nrf2^SMCKO^* mice, with Id2 decreasing and p16 increasing in the *Nrf2^SMCKO^* group (**Figure 6F**). Additionally, a negative correlation between Id2 and p16 expression levels was observed in the RNAseq data (**Figure 6G**). These findings suggest a potential interaction between ID2 and P16 in the context of NRF2’s protective function in VC.

**Figure 6.**
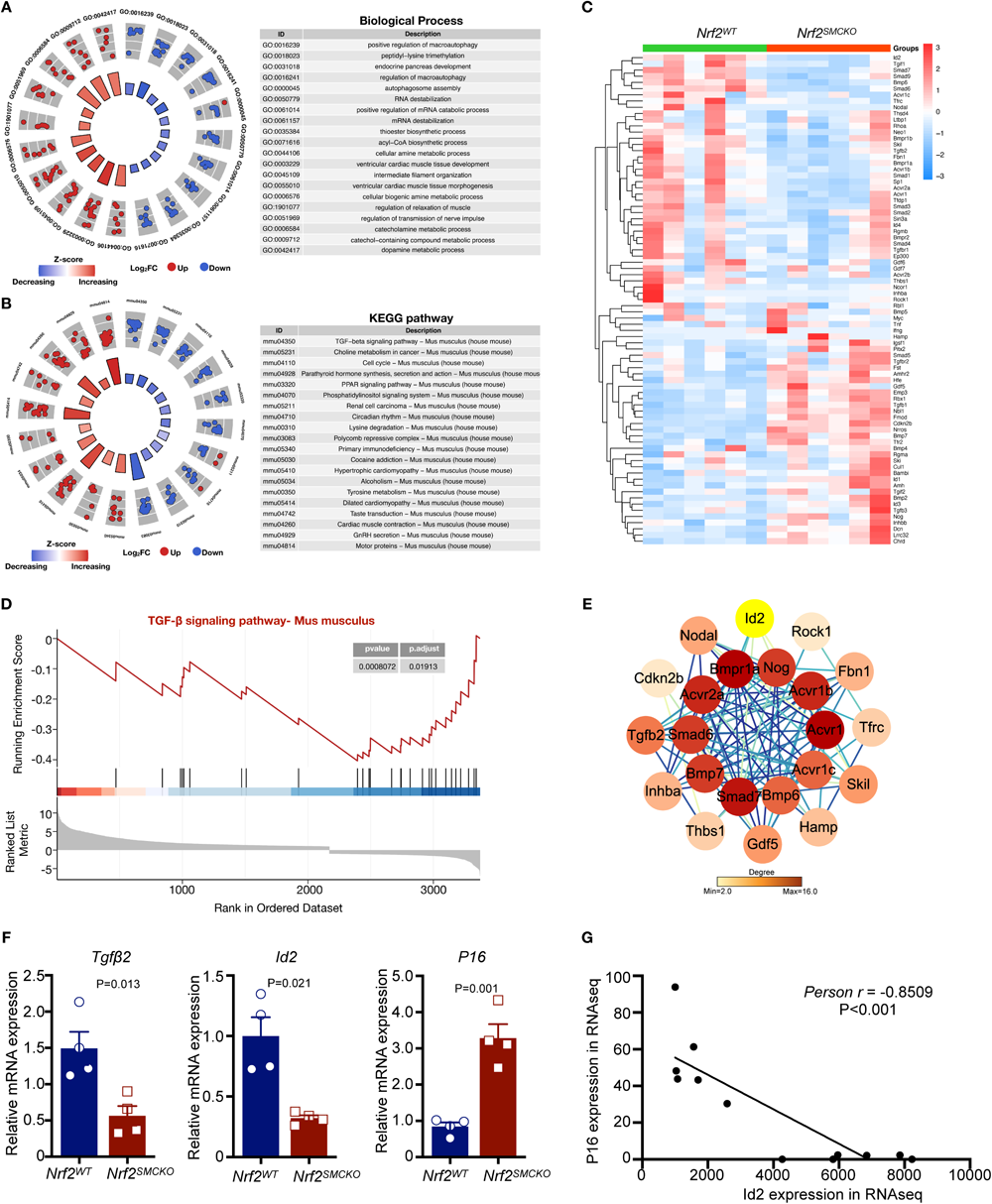
Transcriptional Profiles in *Nrf2*^SMCKO^ Mice Aorta. **A-B,** The top 20 significantly enriched Gene Ontology Biological Processes (GO-BP) categories and KEGG pathways from differentially expressed genes (DEGs) between *Nrf2^SMCKO^* and *Nrf2^WT^* mouse aortas. **C,** Heatmap illustrating DEGs enriched in the TGFβ signaling pathway. **D,** Gene Set Enrichment Analysis (GSEA) of *Nrf2^SMCKO^* and *Nrf2^WT^* mouse aortas. **E,** Protein-Protein Interactions (PPI) analysis of DEGs involved in the TGFβ signaling pathway. **F,** Quantitative real-time PCR analysis of Id2 and p16 in *Nrf2^SMCKO^* and *Nrf2^WT^*mouse aortas (n=4 per group). **G,** Correlation analysis between Id2 and p16 expression using RNA-seq data. All data are presented as mean ± SE. Statistical significance was determined using a 2-tailed unpaired Student t-test or 1-way ANOVA with Tukey’s multiple comparisons test.

### ID2 Contributes to the Protective Effect of NRF2 in VC

To investigate the potential link between ID2 and NRF2’s beneficial effects in VC, we revisited both the Vitamin D-mediated VC mouse model and the high-Pi-induced VSMC calcification model. In *Nrf2^SMCKO^*mice aortas, Id2 nuclear translocation was significantly diminished compared to *Nrf2^WT^* counterparts. Vitamin D treatment exacerbated this reduction in Id2 nuclear translocation *in vivo* (**Figure 7A**). IF staining revealed that vitamin D treatment increased p16 expression, with Nrf2 deficiency leading to a further rise in P16 protein levels (**Figure 7B**). *In vitro,* while high-Pi diminished Id2 expression, induced Nrf2 overexpression via LV-NRF2 notably elevated Id2 expression at both mRNA and protein levels compared to LV-CON (**Figures 7C** and **7E**). Conversely, TGFβ1 and p-Smad3 protein expressions were augmented under high-Pi conditions, but were significantly diminished by Nrf2 overexpression (**Figures 7C**, **7E,** and **7F**). Notably, high-Pi increased VSMC senescence, as indicated by a rise in SA-β gal positive cells, an effect that was mitigated by Nrf2 overexpression (**Figure 7D**). Furthermore, qPCR analysis showed that high-Pi induced an upsurge in cell senescence markers (p16 and p21) and SASP components (Il-6, Il-8, Ccl2, Cxcl1, Csf2, Opg). Remarkably, Nrf2 overexpression reduced high-Pi-induced cell senescence, as shown by lower levels of senescence markers (p16 and p21) and SASP components (Il-6, Il-8, Cxcl1, Csf2, Opg) (**Figure 7G**), underscoring the potent role of NRF2 in reducing VSMC calcification and cellular senescence.

**Figure 7.**
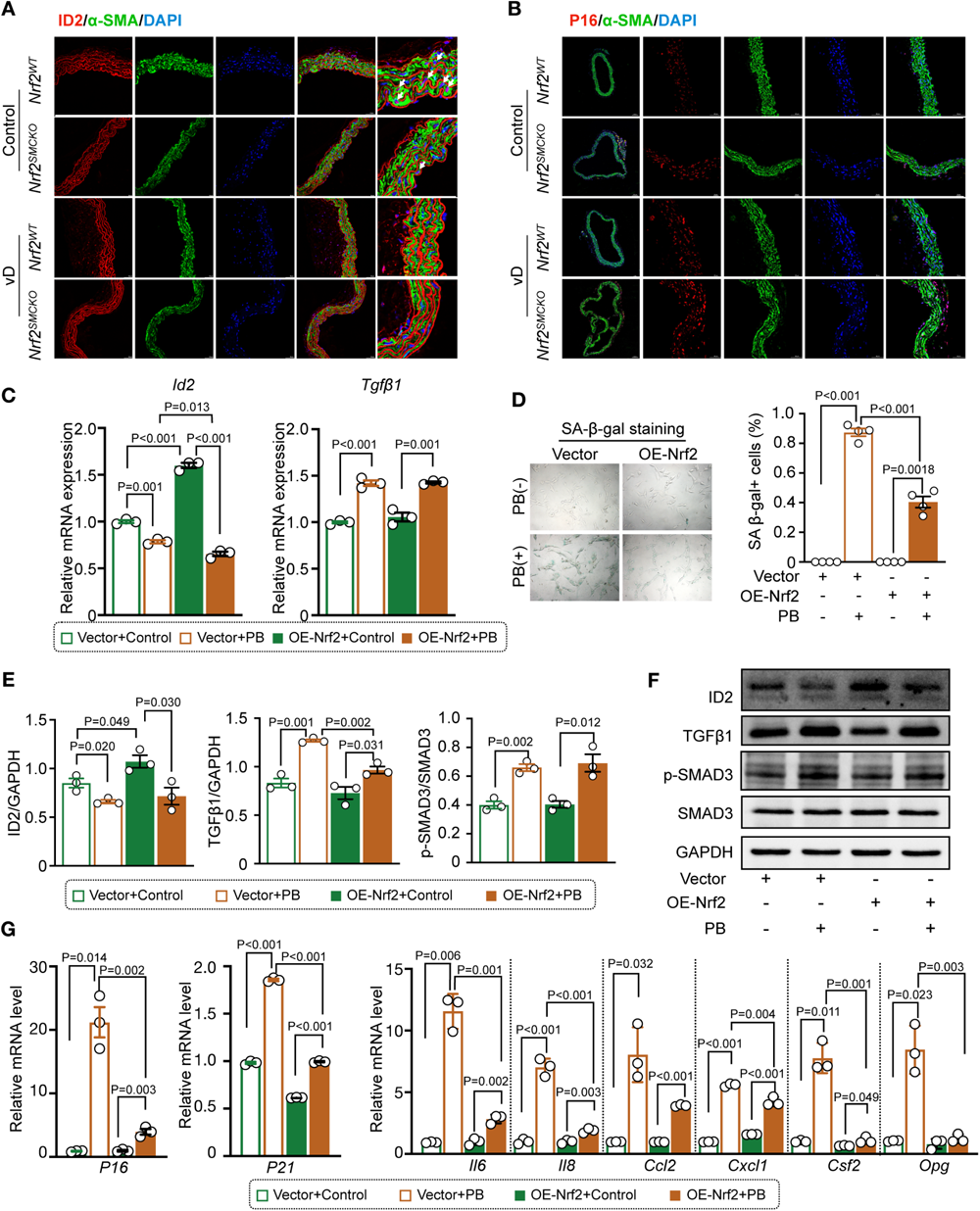
Nrf2 Overexpression Upregulated Id2 and Alleviated VSMC Senescence. **A,** Immunofluorescence staining for Id2 (red) and α-SMA (green) in serial sections of *Nrf2^SMCKO^* and *Nrf2^WT^* mouse aortas following injection with dextrose water or vitamin D. Arrows indicate areas of intranuclear Id2 upregulation. **B,** Immunofluorescence staining for P16 (red) and α-SMA (green) in serial sections of *Nrf2^SMCKO^* and *Nrf2^WT^*mouse aortas injected with dextrose water or vitamin D. **C,** Quantitative real-time PCR analysis of Id2 and Tgfβ1 in MOVAS cells treated with NRF2 lentivirus or control lentivirus, with or without 2.6 mmol/L High-Pi stimulation (n=3 per group). **D,** Representative micrographs of SA-β-gal staining and quantification of the percentage of SA-β-gal positive cells in MOVAS cells treated with NRF2 lentivirus or control lentivirus under High-Pi stimulation (n=4 per group). **E,** Western blot analysis of Id2, Tgfβ1, p-Smad3, and Smad3 in MOVAS cells treated with NRF2 lentivirus or control lentivirus, with or without High-Pi stimulation (n=3 per group). **F,** Quantitative real-time PCR analysis of cell senescence markers (p16 and p21) and SASP components (Il-6, Il-8, Ccl2, Cxcl1, Csf2, Opg) in MOVAS cells treated with NRF2 lentivirus or control lentivirus under High-Pi stimulation (n=3 per group). Scale bars represent 100μm and 50μm. All data are presented as mean ± SE. Statistical significance was determined using a 2-tailed unpaired Student t-test or 1-way ANOVA with Tukey’s multiple comparisons test.

To further clarify Id2’s role in Nrf2-mediated regulation of VSMC mineralization, Id2 was targeted using specific siRNA. The successful downregulation of Id2 was verified at both mRNA and protein levels (**Figures 8A** and **8B**). Under high-Pi conditions, Nrf2 overexpression significantly curtailed calcium deposition and the Alizarin red-positive area (**Figures 8C** and **8D**), along with a reduction in *Bmp2* and *Runx2* mRNA expression in VSMCs (**Figure 8E**). However, Id2 knockdown compromised these Nrf2-mediated protective effects (**Figures 8C**, **8D**, and **8E**). Under high-Pi conditions, *Id2* silencing undermined NRF2’s efficacy in diminishing oxidative stress, as evidenced by DHE staining (**Figure 8F**), and lessened Nrf2’s capacity to mitigate DNA damage, as shown by γ-H_2_AX IF staining (**Figure 8F**). Id2 was also found to be instrumental in Nrf2’s suppression of high-Pi-induced cellular senescence, indicated by SA-β gal staining (**Figures 8G and 8H**). qPCR analysis revealed that while Nrf2 overexpression led to a decrease in senescence markers (p16 and p21) and SASP components (Il-6, Ccl2, Cxcl1, Csf2, Opg) under high-Pi conditions. Id2 silencing increased these senescence markers and reversed the beneficial modulatory effects of NRF2 on p16, p21, and certain SASP components (Il-6, Ccl2, Cxcl1, Csf2, Opg) (**Figure 8I**). These findings suggest a critical interaction between ID2 and NRF2 in modulating VC, implicating a significant role of P16-related cellular senescence in this process.

**Figure 8.**
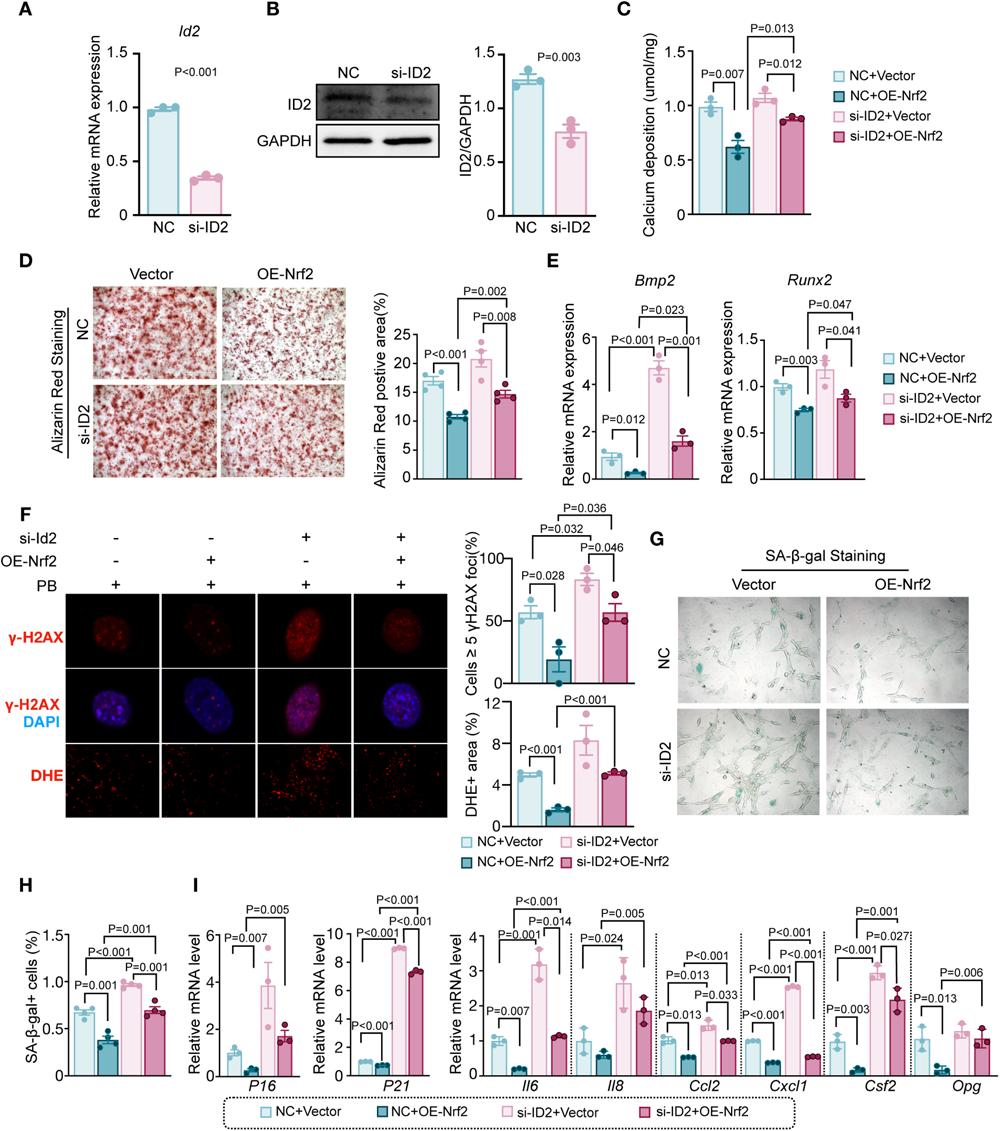
ID2 Contributes to the Protective Effect of NRF2 in VC. **A-B,** Quantitative real-time PCR and Western blot analyses assess Id2 expression in MOVAS cells treated with Id2 siRNA or control siRNA (n=3 per group). **C,** Quantitative analysis of calcium deposition in MOVAS cells treated with various combinations of control siRNA, NRF2 lentivirus, and Id2 siRNA under 2.6 mmol/L High-Pi stimulation (n=3 per group). **D,** Representative micrographs of Alizarin Red S staining and quantification of the percentage of Alizarin Red S-positive area under each treatment condition (n=4 per group). **E,** Quantitative real-time PCR analysis of Bmp2 and Runx2 under each treatment condition (n=3 per group). **F,** Representative micrographs of γ-H2AX and DHE staining under each condition, alongside quantification of the percentage of cells with 5 or more γ-H2AX foci and DHE-positive areas (n=3 per group). **G-H,** Representative micrographs of SA-β-gal staining and quantification of the percentage of SA-β-gal positive cells under each treatment condition (n=4 per group). **I,** Quantitative real-time PCR analysis of cell senescence markers (p16 and p21) and SASP components (Il-6, Il-8, Ccl2, Cxcl1, Csf2, Opg) under each treatment condition (n=3 per group). Scale bar represents 20μm. All data are presented as mean ± SE. Statistical significance was determined using a 2-tailed unpaired Student t-test or 1-way ANOVA with Tukey’s multiple comparisons test.

## DISCUSSION

VC is a complex phenomenon commonly observed in individuals with chronic kidney disease, atherosclerosis, diabetes, and advancing age^36^. It significantly contributes to the pathophysiological progression of diseases by increasing arterial stiffness and consequently elevating the risk of cardiovascular morbidity and mortality^37^. OS is recognized as a pivotal regulator in this process, affecting VC through cell differentiation, apoptosis, senescence, inflammation, and extracellular matrix remodeling^6^. However, the precise mechanisms by which NRF2 influences the progression of VC and how NRF2 establishes a connection between aging and VC remain elusive. Our study revealed that aging accelerates VC in mice, correlating with diminished NRF2 activity, increased OS, and reduced arterial apoptosis, thereby underscoring the profound connection between aging and VC. Moreover, specific knockout of Nrf2 in VSMCs aggravated VC across various calcification models, whereas overexpression of Nrf2 curtailed calcium deposition by attenuating VSMC senescence and apoptosis. Age-associated NRF2 dysfunction possibly brought on VC by accelerating VSMC senescence. Through RNA-seq analysis of aortas in *Nrf2*^SMCKO^ and control mice, we have identified a novel downstream target of NRF2, namely ID2, which is indispensable for the protective effect of NRF2 in VC. In conclusion, our study shed light on the accelerated calcification observed in the context of aging. The protective effects of Nrf2 overexpression in VSMC and identifying ID2 as a downstream target contribute to our understanding of the complex mechanisms underlying VC. Further research is warranted to develop novel therapeutic Interventions aimed at mitigating the burden of VC-associated complications.

Evidence is increasingly suggesting that activation of NRF2 by various pharmacological agents and natural compounds can prevent VC in both in vitro and in vivo settings^13–17,22^. However, the function of NRF2 activators lacks specificity and may induce activation of alternative pathways, such as the NF-κB signaling pathway, which has also been implicated in the inhibition of VC. In vitro, silencing *Nrf2* or accelerating degradation of NRF2 have also been proven to promote VSMC calcification induced by high phosphate. In turn, overexpressing *Nrf2* or retarding degradation of NRF2 inhibits VSMC calcification^16,21,23,38^. Similarly, our study demonstrates that the overexpression of *Nrf2* reduces the expression of Bmp2 and inhibits calcium deposition in VSMC, accompanied by a decrease in cell senescence and apoptosis. It is essential, however, to acknowledge that NRF2 activators’ effects are not confined to a single organ or tissue and vary across different cell types. The explicit role of NRF2 in vascular smooth muscle tissue *in vivo* remains relatively unexplored. To address this knowledge gap, we conducted a VSMC-specific knockout of *Nrf2* transgenic mice on a C57BL/6J background and demonstrated the protective effect of NRF2 in a high adenine diet-induced and Vitamin D3-induced calcification model. The aorta of *Nrf2^SMCKO^* mice exhibited a more pronounced calcification than the control group, with attenuating VSMC senescence and apoptosis. Additionally, our *ex vivo* aortic ring culture experiments aligned with these obeserevations. Furthermore, both primary VSMCs derived from *Nrf2^SMCKO^* mice and VSMC cell lines transfencted with Nrf2 siRNA displayed significant calcification under high-phosphate conditions *in vitro*, providing robust evidence of NRF2’s anti-calcification influence in VSMCs.

Imbalances between oxidative and antioxidant systems are a notable consequence of aging, significantly influencing the progression of various chronic diseases such as CVD, diabetes, neurological disorders, and CKD^39^. As a master regulator in the antioxidant response, NRF2 is capable of activating numerous tissue-specific cytoprotective proteins throughout an organism’s lifespan^40,41^. Nevertheless, the aging process impairs the inducible activity of NRF2 in the aortas of monkeys and rats, leading to an increase in ROS production^28,29^. Our investigations have revealed that aging diminishes the expression and activity of NRF2 in arterial tissues of mice, thereby exacerbating OS. This underscores the crucial role of age-associated NRF2 dysfunction in the imbalance between oxidants and antioxidants. Arterial calcium levels and cardiovascular disease prevalence are known to increase with age^25^. Yet, most studies exploring the relationship between aging and VC have been observational, lacking experimental evidence for a comprehensive understanding. Utilizing a mouse model with both young and aged cohorts, our study has demonstrated that aging accelerates VC, associated with increased OS and reduced apoptosis. We also confirmed that NRF2 activators, such as DMF and tBHQ, can reverse VC in aged mouse aortas, positioning NRF2 as a key mediator in the nexus between aging and VC. Aging is associated with an accumulation of senescent cells resistant to apoptosis, contributing to medial calcification and increased expression of osteogenic markers^7,42^. Silencing Nrf2 has been found to speed up the senescence of vascular cells^43,44^, with both senescent cell accumulation and Nrf2 dysregulation implicated in medial VC pathogenesis^20^. The burgeoning body of evidence suggests that activating NRF2 could provide protective effects against age-related VC by mitigating cellular senescence.

In a quest to explore the target genes of NRF2 in VC, we embarked on an RNA-seq analysis of aortas in *Nrf2^SMCKO^* and *Nrf2^WT^* mice. The functional enrichment analyses of DEGs have indicated that the downregulated genes in *Nrf2^SMCKO^* mice aortas were primarily enriched in the TGFβ signaling pathway. The TGFβ superfamily, including TGF-βs, BMPs, and activins, is known to profoundly influence the initiation and progression of VC. Elevated TGFβ levels, along with specific osteogenic BMPs like BMP2 and BMP4, are commonly seen in early medial calcific lesions, especially in CKD-induced VC^45^. On the other hand, BMP7 emerges as a critical inhibitor of VC^46^. In our study, PPI analysis has identified ID2 as the core of DEGs enriched in the TGFβ signaling pathway. ID2, part of the DNA-binding protein inhibitor family, plays a crucial role in cell proliferation and differentiation. It interacts with the retinoblastoma protein (pRb), neutralizing its growth-suppressing function and modulating cell differentiation. This interaction is particularly noteworthy given that ID2, but not ID1 or ID3, can counteract the cell cycle arrest imposed by p16, a known selective inhibitor of pRb and related kinases^47^. ID2 also promotes a pro-survival and proliferative state in VSMCs in response to various stressors^48^, underscoring its significance in cellular adaptation. Interestingly, ID2 responds to TGF-β/BMP signaling^49^, influencing VSMC phenotype transitions from contractile to synthetic forms through the PI3K/AKT/ID2 pathway^50^. The BMP-SMAD-ID signaling cascade has been shown to counteract p16^INK4A^-mediated cell senescence, indicating a protective role against cellular aging^51^. Remarkably, ID2 has been identified as a pivotal factor in osteogenic differentiation. Prior research has shown that ID2 suppresses osteogenesis in C2C12 cells by inhibiting the activation of core binding factor α-1^52^. Notably, utilizing an Id2 knockout mouse model has revealed that ID2 promotes cartilage formation by enhancing BMP signals through the reduction of Smad7 expression^53^. Additionally, ID2 is known to curtail ROS production caused by glucose deprivation, alleviate mitochondrial damage, and decrease cell death due to calcium overload^54^. In our studies, we noted a decrease in Id2 expression in *Nrf2^SMCKO^* mouse aortas. Conversely, overexpressing Nrf2 led to an upregulation of Id2. The absence of ID2 mitigated the NRF2-mediated inhibition of VSMC calcification. Moreover, ID2 participated in the NRF2 regulatory pathway during high-phosphate-induced VSMC senescence. Correspondingly, our research showed a decline in Nrf2 expression in ID2-deficient VSMCs. Earlier studies have demonstrated that ID2 activates NRF2 to protect retinal pigment epithelium cells from oxidative damage^55^. These findings highlight the intricate interplay between ID2 and NRF2 in managing oxidative stress and cell senescence, underscoring their significant roles in vascular calcification.

In conclusion, our research delves into the critical role of NRF2 in shielding VSMCs from calcification by counteracting OS and cellular senescence through activating ID2. The underlying molecular mechanisms present significant potential in combating VC, a major contributor to cardiovascular diseases. Intriguingly, the dynamic interaction between NRF2 and its downstream pathways opens up new avenues for understanding the relationship between aging and VC. However, given the intricate interplay of these pathways, further research is essential to comprehend the detailed molecular regulation within the NRF2-ID2 axis fully. It is through these detailed studies that we can anticipate developing therapeutic strategies against VC and age-related cardiovascular diseases.

### Nonstandard Abbreviations and Acronyms

VC: vascular calcification OS: oxidative stress
NRF2: nuclear factor erythroid 2-related factor 2 ID2: inhibitor of DNA binding 2
VSMC: vascular smooth muscle cell CKD: chronic kidney disease
CRF: chronic renal failure ESRD: end-stage renal disease
SASP: senescence-associated secretory phenotype ROS: reactive oxygen species
SOD: superoxide dismutase

## Acknowledgments

The authors thank Pennington Biomedical Research Center/LSU System for presenting *Nrf2^f^*^lox/flox^ mice.

## Supplemental Material

Figure S1-S4 Tables S1-S2

**Figure S1. Oxidative Stress and NRF2 Nuclear Translocation in Aged Mice Aortas**

**A,** Representative micrographs of DHE staining, along with quantification of the percentage of DHE-positive areas in serial sections of aged and young mice aortas, both with and without 3.8 mmol/L High-Pi stimulation (n=3 per group). **B,** Immunofluorescence staining of NRF2 in serial sections of aged and young mice aortas under High-Pi stimulation. Scale bars represent 100μm and 50μm. All data are presented as mean ± SE. Statistical significance was determined using a 2-tailed unpaired Student t-test or 1-way ANOVA with Tukey’s multiple comparisons test.

**Figure S2. PCR genotyping and Western blot analysis of NRF2 for *Nrf2^SMCKO^* mice**

**A,** PCR genotyping of genomic DNA from *Nrf2^SMCKO^* (*Tagln-Cre^/+^; Nrf2^flox/flox^*) and *Nrf2^WT^*(*Tagln-Cre^/-^; Nrf2^flox/flox^*) mice to confirm the specific knockout. **B,** Western blot analysis of NRF2 protein levels in the heart, liver, skeletal muscle, and uterus of *Nrf2^SMCKO^* and *Nrf2^WT^* mice (n=3 per group). All data are presented as mean ± SE. Statistical significance was determined using a 2-tailed unpaired Student t-test or 1-way ANOVA with Tukey’s multiple comparisons test.

**Figure S3. Nrf2 silencing Irritated VSMC Calcification**

**A-B,** Quantitative real-time PCR and Western blot analyses assess the expression of Nrf2 and Nqo1 in MOVAS cells treated with Nrf2 siRNA or control siRNA (n=3 per group). **C,** Representative micrographs of Alizarin Red S staining, encompassing both whole well and microscopic views (scale bar, 20μm), along with quantification of the percentage of Alizarin Red S-positive area in MOVAS cells treated with Nrf2 siRNA or control siRNA, both with and without High-Pi stimulation (n=3 per group). **D,** Quantitative analysis of calcium deposition in MOVAS cells treated with Nrf2 siRNA or control siRNA, under both conditions (n=3 per group). Scale bar represents 20μm. All data are presented as mean ± SE. Statistical significance was determined using a 2-tailed unpaired Student t-test or 1-way ANOVA with Tukey’s multiple comparisons test.

**Figure S4. Nrf2 Overexpression reduced VSMC apoptosis**

**A,** Detection of cell apoptosis using Annexin V-FITC/PI in MOVAS cells treated with NRF2 lentivirus or control lentivirus, under both normal and High-Pi conditions (n=3 per group). **B,** Western blot analysis of cleaved-Caspase3 and Caspase3 in MOVAS cells treated with NRF2 lentivirus or control lentivirus, with or without High-Pi stimulation (n=3 per group). All data are presented as mean ± SE. Statistical significance was determined using a 2-tailed unpaired Student t-test or 1-way ANOVA with Tukey’s multiple comparisons test.

**Table S1. Primers applied in real-time PCR**

**Table S2. Differentially expressed genes (DEGs) identified in transcriptional profiles**

